# Genomic Characterization and New Species Identification of *Winogradskyella* with Comparative Analysis of Polysaccharide Utilization Loci

**DOI:** 10.1101/2025.03.31.646394

**Authors:** An-Qi Sun, Jia-Yu Deng-Li, Han-Zhe Zhang, Jin-Hao Teng, Zong-Jun Du, De-Chen Lu

## Abstract

Marine macroalgae convert a substantial fraction of fixed carbon dioxide into various polysaccharides. *Winogradskyella* is a genus within the phylum *Bacteroidota* with a clear marine origin especially from marine macroalgae. Most members of this genus have been found associated with macroalgae or microalgae. Using a combination of genomic, phylogenetic, and biochemical approaches, we characterized five novel *Winogradskyella* species capable of degrading marine macroalgal polysaccharides. Bioinformatic PUL annotations suggest usage of a large array of polysaccharides, including laminarin, α-glucans, and alginate as well as mannose- and fucose-, highlighting the genus’ involvement in the marine carbon cycle. Many of the PULs exhibit new genetic architectures and suggest substrates rarely described for marine environments. Using the SusC/D trees, we analyzed evolutionary relationships of SusC/D homologs and found indications for profound changes in microbial utilization of laminarin, alginate, α-glucans, β-mannan, and sulfated xylan. Through the genomic study and annotation of PULs, we explored the complex relationship between genome size and polysaccharide degradation potential of *Winogradskyella*. These findings underscore the ecological significance of *Winogradskyella* in marine ecosystems and their role in polysaccharide degradation, providing a basis for biogeochemical cycling and future biotechnological applications.

## Introduction

The omnipresence of microorganisms stems from their remarkable metabolic adaptability, enabling them to thrive in virtually every conceivable habitat on Earth. Most habitats host unknown microbial populations, and macroalgal epiphytic environments represent a clear example. The surface of algae hosts a dynamic and intricate micro-environment, shaped by the colonization of various planktonic microorganisms, including bacteria, fungi, diatoms, and other organisms. (1), and mostly belonging to yet to be described bacterial lineages (2). Physicochemical properties, metabolite composition, defense mechanism (3) and attractant patterns (4), among different groups of macroalgae (*Phaeophyceae, Chlorophyceae and Rhodophyceae*) prompt them to have specific and unique microbial architectures (5). The genus *Winogradskyella* is a monophyletic genus within the *Flavobacteriaeae*, a family of the phylum *Bacteroidota*. The genus was described by Nedashkovskaya et al. based on three species of marine bacteria, *Winogradskyella thalassocola* (type species), *Winogradskyella epiphytica* and *Winogradskyella eximia*(6). At present, genus *Winogradskyella* contained more than 46 species with valid published name listed in LPSN (https://lpsn.dsmz.de/genus/Winogradskyella). Strains of the genus *Winogradskyella* have been discovered in various marine environments, such as sea water (7), plankton (8), marine sediment (9, 10), phycosphere of algae or plants(11, 12), and marine animals (13, 14). The bacteria in the genus *Winogradskyella* are Gram-stain negative and heterotrophic, with yellow-, orange- or salmon-coloured pigmentation. The main cellular fatty acids are straight-chain saturated, branched-chain saturated and unsaturated fatty acids iso-C_15: 0_, anteiso-C_15: 0_, iso-C_15: 1_, iso-C_16:0_ 3-OH and iso-C_17: 0_ 3-OH(15). The polar lipid profile comprises phosphatidylethanolamine and one or two unknown aminolipids(16, 17). The major respiratory quinone is menaquinone MK-6.

In bacteria, the genes for the breakdown and take-up of polysaccharides are often co-located in dedicated polysaccharide utilization loci (PULs), in particular in the *Bacteroidota*. The capacity to degrade various land plant polysaccharides has been well studied in human gut *Bacteroidota* (17), and in some marine *Bacteroidota* targeting algal polysaccharides, e.g., alginate (18), laminarin (19, 20) and carrageenan (20). Polysaccharide Utilization Loci (PULs) are genetic regions in certain bacteria, particularly those in the *Bacteroidota*, that enable the breakdown and utilization of complex polysaccharides. These loci consist of coordinated sets of genes encoding various proteins, including: Carbohydrate-Active Enzymes (CAZymes) include Enzymes like glycoside hydrolases, polysaccharide lyases, and carbohydrate esterases that degrade polysaccharides; Surface Glycan-Binding Proteins (SGBPs): Proteins that recognize and bind specific polysaccharides on the bacterial cell surface; Transport Systems (SusCD complexes): Mechanisms for importing the breakdown products into the cell for metabolism (21). By efficiently degrading and utilizing complex polysaccharides, marine bacteria play a pivotal role in the global carbon cycle, facilitating the recycling of organic carbon in both terrestrial and aquatic ecosystems.

Macroalgae are important primary producers in nearshore ecosystems, significantly contribute to coastal carbon sequestration, ecosystem stability and biodiversity maintenance (22). Pioneering studies on structural elucidation of polysaccharides from macroalgae were performed (23), but precise macroalgal and microalgal polysaccharide structures remain mostly unresolved, because they require sophisticated methods (24). Among these, phycosphere microbiota plays a crucial role in regulating algal growth and transforming carbon, studies have shown a greater synergistic degradation using shared exo-enzymes in phycosphere communities than in the water column (25). Many members of *Bacteroidota*, including marine representatives of the family *Flavobacteriaeae*, are specialized on polysaccharide degradation. PUL analysis of epiphytic bacteria co-occurring with macroalgae could serve as an alternative starting point to advance insight into the structures of marine polysaccharides and to understand their microbial decomposition. Alginate is a polysaccharide, known for its solubility and gel-forming properties. Laminarin, found in red algae, shares structural similarities with alginate but has distinct sugar units. Both polysaccharides play crucial roles in the structure of algal cell walls and act as carbon sources for marine microorganisms. Such as laminarin is a major molecule in the marine carbon cycle (26). Microbes, particularly from the *Bacteroidota*, degrade these polysaccharides, contributing to the carbon cycle in marine ecosystems. Understanding their degradation mechanisms is key to advancing microbial decomposition research and sustainable marine management.

In recent years, several researchers have studied the diversity and function of the epiphytic bacteria associated with macroalgal host (2), but largely these interactions remain underexplored. In the present study we analysed the cultivable diversity of epiphytic bacteria associated with five different marine macroalgae (*Saccharina japonica*, *Grateloupia* sp., *Gelidium* sp. and *Ulva*, sp.) sampled from the Weihai of China. During our survey of polysaccharide utilization capacity of phycosphere microbiota in the coastal zone of Weihai, China, we identified five novel species (A3E31^T^, PE311^T^, 4-2091^T^, 3-1016^T^ and A2^T^) belonging to the genus *Winogradskyella*. Here we present a comparative analysis of PULs from 109 genome of *Winogradskyella* and 21 isolated from the macroalgae, comprising a total of 336 manually determined PULs.

## Methods

### Isolation and culture condition

To investigate the ability of bacteria from the phylum *Bacteroidota* to degrade marine polysaccharides, marine 48 macroalgae, 12 seawater, 12 sediment and a kelp samples were collected from a coastal area in Weihai, China, 37 isolates belonging to the genus *Winogradskyella* were discovered and five belong to novel species. The samples were serially diluted to 10^-5^ in 9 ml sterile seawater, and 100 µl from each dilution was plated on Marine Agar 2216 (MA; Becton Dickinson) plates. These plates were then incubated in an anaerobic environment at 28 °C for one month. Following initial isolation and purification, five distinct colonies were identified: one smooth and convex with an orange pigment, another four exhibiting a yellow-coloured pigment. After identification through 16S rRNA gene sequence analysis and repeated streaking for purification, the purified colonies were routinely maintained at 28 °C on MA for one week. The strains were preserved at -80 °C in a solution containing 3.0% salt and 20.0% (v/v) glycerol. The reference strains *Winogradskyella* species, we acquired *Winogradskyella thalassocola* KCTC 12221^T^, *Winogradskyella aquimaris* KCTC 23502^T^ and *Winogradskyella echinorum* KCTC 22026^T^ were obtained from the Korean Collection for Type Cultures (KCTC).

### Physiological characterization

Phenotypic characterizations were performed during the exponential period of growth. Accurate verification of negative or positive results was performed according to the bioMérieux Gram staining kit manual. Colony surface morphology was observed using a light microscope (E600, Nikon) and a scanning electron microscope (FEI Nanonova SEM450) (27). Gliding motility was examined in modified marine broth 2216 (MB; BD) supplemented with 0.3% agar as described previously (28). The range of temperature for growth was tested at the following temperatures: 4, 15, 20, 25, 28, 30, 33, 35, 37 and 40 °C. The pH was modulated to pH 5.5–9.5 (0.5 pH units apart) by adding the same densities and dissimilar pH value buffers in marine broth 2216 (MB; Becton Dickinson) to detect the effect of pH on cellular development (29). Bacteria were cultivated in derivative MA supplemented with artificial seawater without NaCl as the basal medium, and their salinity and salt tolerance were evaluated. The salt gradient of the medium was set at 0–10.0% NaCl (0.5 % intervals). All experiments were repeated three times unless otherwise noted. MA with or without 0.1 % (w/v) KNO_3_ was placed together in anaerobic jars and incubated in anaerobic and microaerobic environments for 2 weeks to observe growth (30).

Testing for catalase activity with a 3% H_2_O_2_ solution and oxidase activity using an oxidase test kit from bioMérieux. Anaerobic growth was assessed over 15 days at 30°C in an anaerobic bag, on modified MA medium with or without 0.1% (w/v) NaNO_3_. Hydrolytic capabilities for agar, starch, alginate, casein, CM-cellulose, DNA, and lipases (Tweens 20, 40, 60, and 80) were evaluated according to the CLSI (2021) method. API 20E and API 50CH tests were conducted in accordance with the manufacturer’s instructions, except the salinity was adjusted to 3%. Enzyme activities were assessed using the API ZYM test from bioMérieux. Oxidation of various compounds was tested in Biolog GEN III microplates, following the manufacturer’s guidelines. All API and Biolog tests were performed with three biological replicates and two reference strains each time.

### Chemotaxonomic characterization

During the late exponential phase of growth, cell masses from strain A3E31^T^, PE311^T^, 4-2091^T^, 3-1016^T^ and A2^T^, and control strains were harvested from cultures grown on MA under ideal conditions to analyze fatty acids, respiratory quinones, and polar lipids. The harvested cells were freeze-dried, and from these dried cell masses, 10 mg was used to extract fatty acids. These extracts were then analyzed with an Agilent gas chromatograph (model 6890N), and the fatty acids were identified using the Microbial Identification System (MIDI database: TSBA40) (Sasser,1990.). For the analysis of respiratory quinones, 300 mg of the freeze-dried biomasses underwent purification as per the methods described by Minnikin (32) and were subsequently analyzed using high-performance liquid chromatography (HPLC) according to Kroppenstedt et. al (Kroppenstedt, 1982). Polar lipids were extracted using varying ratios of a chloroform/methanol/water solution and analyzed via two-dimensional silica thin-layer chromatography. The identification of total polar lipids was performed using specific detection reagents, following established protocols by Minnikin (32).

### Genome extraction, sequencing, 16S rRNA gene phylogenetic analysis, genomic analysis and annotation of PULs

Genomic DNA of the five strains were extracted and purified using a bacteria genomic DNA kit (Takara). Strains sequencing were performed by Beijing Novogene Biotechnology (Beijing, China) on a NovaSeq (Illumina, San Diego, CA, USA) with 150 bp PE reads at ≥ 100 × coverage. Reads were quality-filtered and assembled with SPAdes v3.9.1 (34) (–careful –cov-cutoff) with k-mer sizes from 27 to 127 bp and a minimum scaffold length of 200 bp. The complete 16S rRNA gene sequences of strain A3E31^T^, PE311^T^, 4-2091^T^, 3-1016^T^ and A2^T^ were extracted from the draft genomes by ContEst16S (www.ezbiocloud.net/tools/contest16s). The returned 16S rRNA gene sequence was submitted to GenBank database and the sequences were analysed using BLAST (https://blast.ncbi.nlm.nih.gov/Blast.cgi) and EzBiocloud (http://www.ezbiocloud.net) to determine their approximate taxonomic affiliations. Phylogenetic trees were reconstructed by the neighbour-joining (NJ) (35), maximum-parsimony (MP) (36) and maximum-likelihood (ML) (37) methods with MEGA 11 (Tamura et al. 2021).

Genes were predicted using Prodigal v2.6.3 (39) and annotated with Prokka. The molecular functions of genes and proteins are associated with ortholog groups and stored in the KEGG orthology (KO) database (40) and the metabolic pathways were analysed in detail employing KEGG’s KofamKOALA server (Kanehisa et al. 2016) (https://www.kegg.jp/blastkoala/, accessed on 3 January 2024). The specific function of gene CAZymes-rich gene in PULs were searched in Carbohydrate Active Enzymes database (http://www.cazy.org/). The prediction and annotation of CAZymes-rich and PULs gene clusters were conducted following the methodology proposed by Dechen et al and R (https://www.r-project.org) generated the related figure. we used dbCAN2 to find these clusters (41).

The presence of gene clusters encoding secondary metabolites was predicted using antiSMASH 6.1.1(42). The DNA G+C content was calculated based on the whole genome sequence. The AAI calculator estimates the average amino acid identity using both best hits (one-way AAI) and reciprocal best hits (two-way AAI) between two genomic datasets of proteins. Average nucleotide identity (ANI) values were calculated using the ChunLab’s online ANI Calculator (43).

### Metagenome-assembled genomes (MAGs), Taxonomic inference of MAGs and abundance analysis of MAGs and draft genome

MAGs from epiphytic belonging to *Ca. Winogradskyella* are part of BioProject PRJEB50838, and have been published and named previously (2, 44). Thirty-eight additional *Winogradskyella* MAGs available in the GTDB database (44). In order to determine MAGs and genomes abundance, metagenomic raw reads were mapped to MAGs using BBMap (minid=99). Reads per kilobase per million (RPKM) was calculated based on MAG length and number of reads mapped.

### Habitat distribution analysis

The 16S high-throughput sequencing data for millions of global samples from multiple habitats were acquired from the Microbial Atlas Project (45) (MAP, https://microbeatlas.org/, accessed 30 July 2024). OTUs that displayed more than 97.0 % 16S rRNA gene sequence similarity to strain A3E31^T^, PE311^T^, 4-2091^T^, 3-1016^T^ and A2^T^ were identified in the genus *Winogradskyella*. The habitat distribution of genus *Winogradskyella* was determined through studying the bacterial abundance of the genus *Winogradskyella* in samples from all over the world.

## Results and discussion

### Characterization of the isolate collection

The 5,527 isolates were assigned to different taxonomic groups, but members of the phyla *Bacteroidota*, *Actinomycetota*, *Pseudomonadota* and *Bacillota* were predominantly isolated (2). Thirty-seven isolates could be assigned to the genus *Winogradskyella* and presumptively identified as phycosphere species, both of which are well-documented inhabitants of the phycosphere across diverse macroalgae species. Five isolates were noval species assigned to the genus *Winogradskyella*. These five isolates were selected for an in-depth taxonomic characterization because of their low 16S rRNA gene sequence similarities to already described species.

### Physiological, phenotypic and chemotaxonomic characteristics of the five selected strains

The five isolates were phenotypically characterized and classified based on their 16S rRNA gene sequences within the genus *Winogradskella*, representing five new species. All strains grew within a temperature range of 27°C to 30°C, and no growth was seen below 4°C or above 40°C. All strains required salt of at least 0.5% NaCl for growth. All isolates were Gram-stain-negative, rod-shaped bacteria and demonstrated aerobic metabolism. The phenotypic attributes distinguishing these five isolates from their closest phylogenetic counterparts are detailed in supplementary Table S1. A combination of physiological tests allowed for the differentiation of these strains from their most closely related species. Additional phenotypic traits of the strains are presented in Table S1, which highlights the comparative phenotypic distinction between the five isolates and adjacent strains of *Winogradskella*, including their ability to hydrolyze macromolecular compounds and enzymatic activity.

The chemotaxonomic features of the strains were consistent with those of the genus *Winogradskella*. All strains contained MK-6 as the sole respiratory quinone. The polar lipid profile of strains A3E31^T^, PE311^T^, 4-2091^T^, 3-1016^T^ and A2^T^ consisted of phosphatidylethanolamine, aminolipid, and unidentified lipids (Supplementary Figure S1). Additionally, 3-1016^T^ and A2^T^ also include polysaccharide lyases. The major fatty acid constituents (>10 %) of strains were iso-C_15:0_ and summed feature 3 (C_16:1_ *ω7c*, C_16:1_ *ω6c*). Cellular fatty acid compositions (%) derived from FAME analysis of all five strains and their closely related type strains are shown in Supplementary Table S2.

### High-quality genome sequences of the *Winogradskyella* strains isolated from the macroalgal surface and sediment

The genome size of the strains is around 3.0 – 4.0 Mb with GC content ranging between 31.69% and 38.3%. Among them (Only those with a completeness > 90%), W sp905480385 and *W helgolandensis* Z963^T^ represented the smallest and largest genomes of 2.22 Mb and 4.59 Mb, respectively, as well as the lowest and highest number of predicted genes with 2,923 and 3,622 (Supplementary Table S3). In addition, *W. undariae* Ha325 showed the lowest G + C mol% content with 31.69%, and *W ulvaetocola* 3-1016^T^ the highest with 32.78%. The collection of genomes represented a total of 89 distinct species. Phylogenetic analyses using amino acid sequences from the genomes, conducted with FastTree methods, showed that the five strains do not cluster together as a single group. However, PE311^T^ and 3-1016^T^ show relatively closer clustering, forming a distinct lineage within the *Winogradskyella* (Fig.1). In total, 109 genomes were analyzed. Twenty-one of these were from macroalgal surface and 88 genomes obtained from other locations and available in public repositories (Supplementary Table S3).

**Fig. 1.**
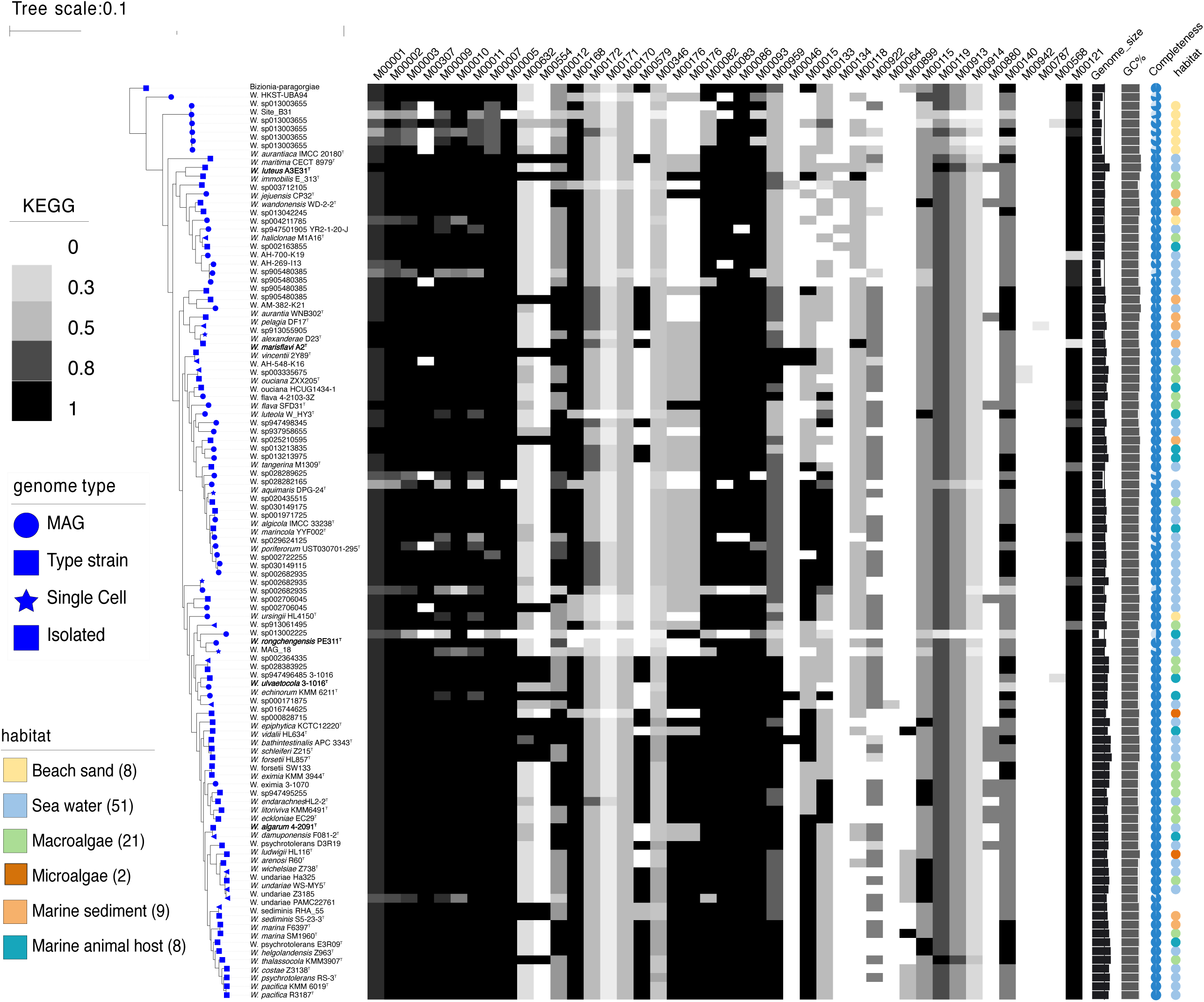
Phylogenetic tree of genomes based on the concatenated alignment of 120 ubiquitous single-copy proteins, highlighting the novel strains A2^T^, A3E31^T^, PE311^T^, 3-1016 ^T^, and 4-2091^T^ within related taxa. Bizionia-paragorgiae was used as the outgroup. The scale bar represents 0.1 substitutions per amino acid position. The isolated habitats, genome sizes, and G+C content of the novel species and their related species are also displayed.

### MAGs retrieved from the macroalgal samples and identified as members of the genus *Winogradskyella*

From the extensive metagenomic data available from macroalgae surface from the years 2022, 1 *Winogradskyella* MAGs was recovered, representing as yet unclassified taxa. The MAGs recovered from *Ulva sp.* (L1_Bin_METABAT 21_1) showed within-group ANI values < 80.06% and AAI values < 80.02%, indicating a strong temporal homogeneity. The MAGs had completeness values >99.45% (Supplementary Table S2), lengths 3.03 Mb and showed the G + C mol% with 33.78%.

We analyzed the genomes of strains isolated from the surface of green and red macroalgae, including strain 3-1016^T^, two strains (3-1070 and 4-2103) belonging to the same genus and a *Winogradskyella* metagenome-assembled genomes (MAGs). Box plot results showed that the RPKM values of all strains was much higher in macroalgae than in seawater and sediment (Fig. 2). This was consistent with the fact that they were isolated from the surface of marine macroalgae. All strains had a significantly higher RPKM values in seawater than sedimen, this suggested that these strains were mainly distributed on the surface of the macroalgae and to a lesser extent in seawater.

**Fig. 2.**
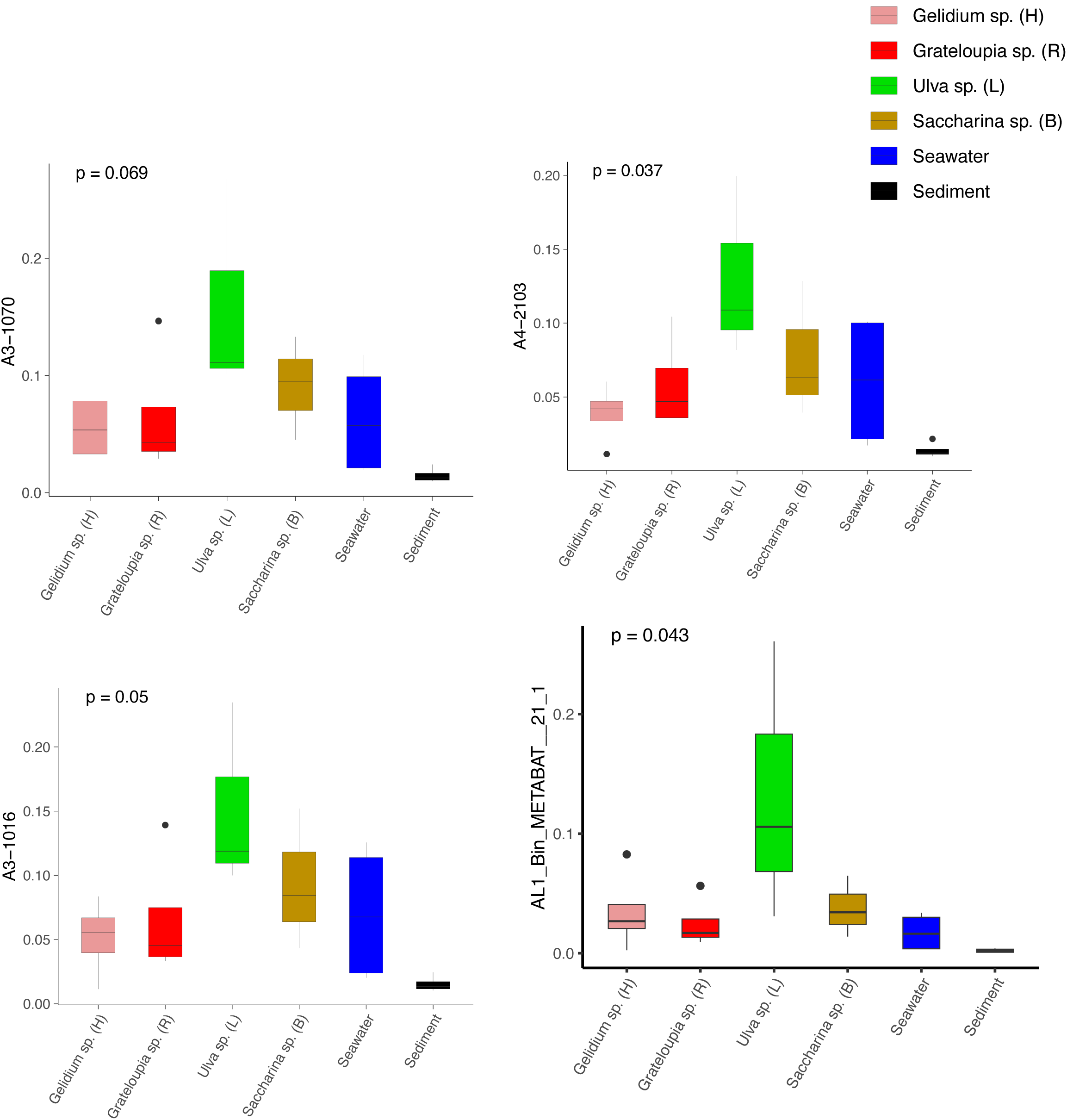
Boxplots show the RPKM values of strains. The data is categorized based on different environmental sources. The p-values corresponding to each comparison are indicated for statistical significance, with a significance threshold of 0.05.

### Phylogenetics, ANI, AAI clustering, phylogenomics, and population genomics

Phylogenetic reconstructions showed consistent topologies regardless of the sequences used to reconstruct them. Specifically, (i) the 16S rRNA genes (Supplementary Fig. S3), (ii) the concatenated sequences of 120 conserved single-copy orthologous genes (essential genes; Fig1). The complete 16S rRNA gene sequences (1526bp, 1526bp, 1526bp, 1522bp and 1526bp) of strains A3E31^T^, PE311^T^, 4-2091^T^, 3-1016^T^ and A2^T^ were obtained. The general features of the genomes are given in Supplementary Table S1. The ANI values between strain A3E31^T^, PE311^T^, 4-2091^T^, 3-1016^T^ and A2^T^ with other strains of the genus were 69.0–79.0%, 68.1-73.2%, 71.3-79.2%, 69.2–80.3%, 72.3–82.9% (Fig. S2), which were far below the standard ANI criteria for species identity (95.0–96.0%). The genome-based phylogenetic analyses were conducted (Fig. 1). The tree indicates close relationships of strains A3E31^T^, PE311^T^, 4-2091^T^, 3-1016^T^ and A2^T^ despite the relatively low level of average branch support. The inconsistencies between the phylogenetic trees based on the 16S rRNA gene sequences and the phylogenomic trees constructed from whole-genome sequence analyses reveal that 16S rRNA gene sequence analyses are insufficient to understand the phylogeny and evolution of the members of the genus *Winogradskyella*. Both the sequence similarities and phylogenetic relationships indicated that strains A3E31^T^, PE311^T^, 4-2091^T^, 3-1016^T^ and A2^T^ represent five novel species of the genus *Winogradskyella*.

The phylogenomics tree can be roughly divided into 4 branches. The first branch is relatively small and contains fewer type strains, but it includes a higher number of MAG. This suggests a more heterogeneous group of strains, with some strains possibly being less well-studied. The second branch is also small in size, containing seven strains compared to other branches, and exhibits moderate genomic variation. The fourth branch, however, is the largest and includes the most model species. It represents a highly diverse group in terms of genomic size and ecological diversity, with strains from various environments. Overall, this division provides valuable insights into the evolutionary, ecological, and genomic diversity among these strains, highlighting the differences in model species representation and the variability in genomic characteristics across various habitats. A detailed summary of the overall genome-relatedness indices is provided in Table S2.

### Habitat distribution analysis

Members of the genus *Winogradskyella*, having been isolated from a myriad of environments, are known for their widespread ecological distribution as exemplified by the studies reporting the isolation of flavobacteria from soil, freshwater, algae and fish (46). And from our study, it can be seen that the five strains were most distributed in aquatic environments, significantly more than in soil and plants. A2^T^ exhibits the highest abundance in marine environments, with a significant value of 16.99%. In contrast, A3E31^T^ shows a moderate abundance in marine environments (0.43%). The abundance of A3E31^T^ in the environment is very low compared to the other four species.

These variations indicate that the five species have distinct habitat distributions, with some preferring marine environments (such as A2 ^T^, which has the highest abundance in marine habitats) and others showing more balanced distributions across various environments. The species A2^T^ stands out with the highest abundance in marine samples, while 3-1016 ^T^ also shows significant presence in marine habitats. On the other hand, species like A3E31^T^ have a more dispersed distribution, with notable abundance in both marine and sediment samples. These observations highlight the ecological diversity and niche specialization of the species under study. This distribution is consistent with the fact that they were isolated from macroalgae living in the ocean (Fig. 3). The radar plots further demonstrate the dominant presence of these strains in aquatic habitats, especially marine environments, which is reflective of their origin from marine algae. Notably, there was minimal distribution in soil or plant habitats, aligning with their ecological characteristics and environmental preferences.

**Fig. 3.**
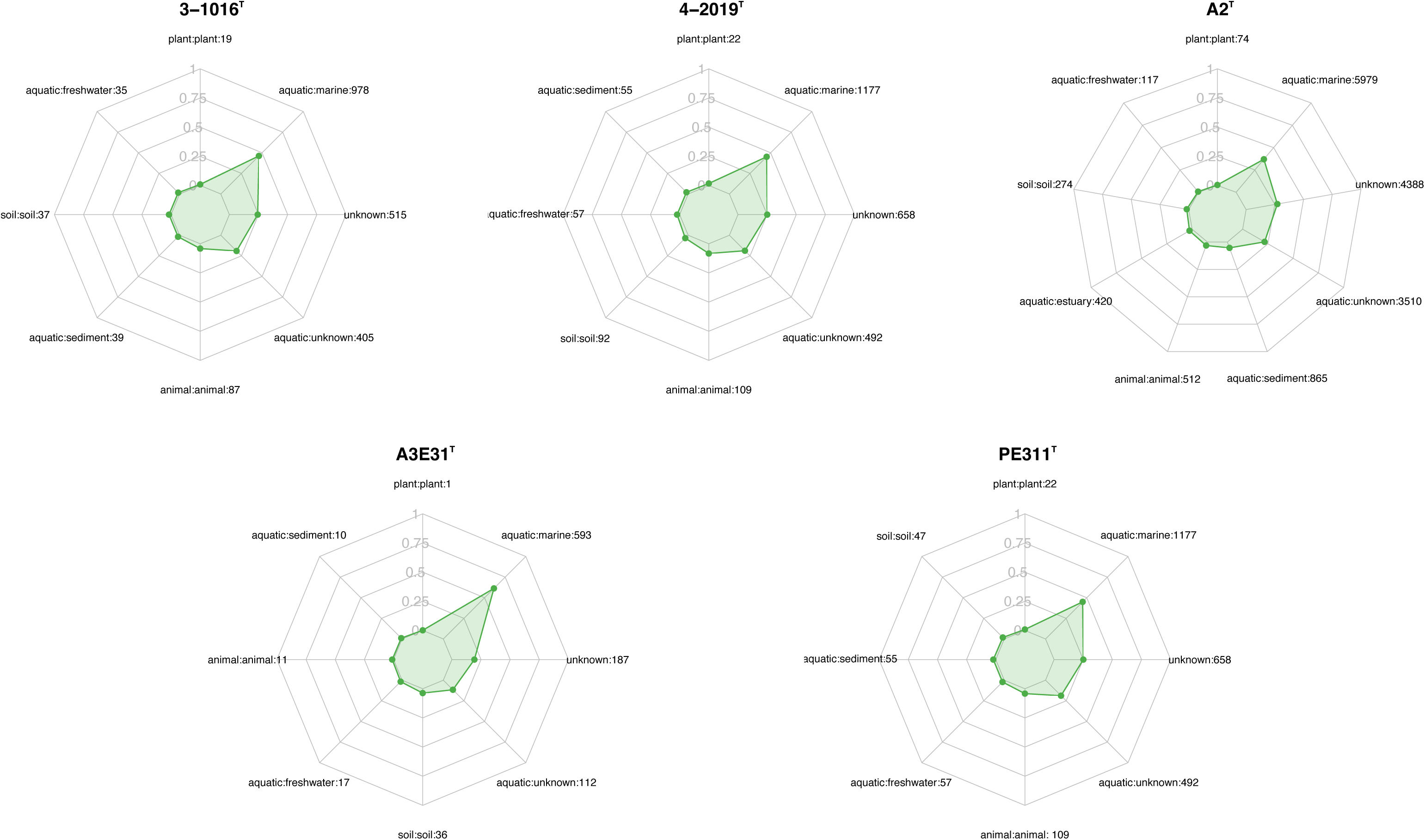
Ecological distribution of *Winogradskyella* strains across various habitats, as shown in the radar plots for different strains (A2^T^, A3E31^T^, PE311^T^, 3-1016^T^, and 4-2091^T^). Each plot illustrates the relative abundance of the strains across multiple environmental categories, including plant, aquatic freshwater, aquatic marine, aquatic sediment, animal habitats, aquatic unknown, soil, and unknown habitats. The numbers next to each habitat label represent the number of sequences identified within that category. The green shaded area in each radar plot indicates the strain’s distribution in that habitat, with the scale ranging from 0 to 1.

### Metabolic pathways

Concerning central carbon metabolism, complete or almost complete glycolysis and gluconeogenesis were found in most members from natural and host-related environments, like the first described species of *Winogradskyella* in this study (Fig.1), suggesting that *Winogradskyella* was a chemoorganotrophic group and may adapt to growing conditions where carbon sources are absent or limited by converting non-sugar substances. Such as Glycolysis (M00001), A3E31^T^, PE311^T^, 4-2091^T^ and 3-1016^T^ show value of 0.8889 and A2^T^ reached 1, respectively, indicating significant metabolic activity in this pathway. This suggests that these strains may efficiently acquire energy through glycolysis in environments with limited carbon sources. Additionally, the gluconeogenesis pathway (M00003) also shows high activity in these strains, especially A3E31^T^ and PE311^T^. The values of these strains are close to 1, indicating their ability to synthesize glucose through gluconeogenesis to support growth.(47)

Lipid metabolism involves the breakdown, synthesis, and storage of lipid, with fatty acids playing a crucial role within this process. The initiation and elongation steps of fatty acid synthesis were observed in the majority of *Winogradskyella* members (Fig. 1). Amino acids also play important roles in bacterial nutrition and growth as essential energy sources and fundamental building blocks of the proteome. Although certain amino acid syntheses remained complete in most MAGs from a particular habitat, the majority of *Winogradskyella* members across various environments exhibited Polyamine biosynthesis auxotrophies (M00134), indicating their dependence on external sources of polyamines for growth (Table S6).

And the presence of complete heme biosynthetic pathways (M00121) in these five bacterial strains suggests that they possess the necessary machinery to autonomously synthesize heme, a vital cofactor for many enzymes involved in cellular respiration and detoxification processes.(Fig.S4) Heme acts as a central component in the electron transport chain, supporting aerobic metabolism and oxidative stress resistance, which are crucial for survival and adaptation to various environmental conditions (48). The ability to independently produce heme reduces reliance on external sources, conferring ecological advantages, especially in nutrient-limited environments. Additionally, Zhang et al. (2024) demonstrated that many marine bacteria rely on heme as a growth-promoting factor, often depending on microbial interactions for heme acquisition(49). However, the presence of a complete heme pathway in these strains indicates they do not require such interactions, enhancing their metabolic flexibility and ecological independence. These strains possess a complete heme biosynthetic pathway, allowing them to independently synthesize heme. Despite this, they are still capable of providing modified compounds to other microbial groups within the biofilm. This ability to function autonomously not only supports their survival in diverse environments but also highlights their potential involvement in biogeochemical cycles, particularly those related to iron.

### Prediction of secondary metabolites

The genomes of the five strains (A2^T^, A3E31^T^, PE311^T^, 3-1016 ^T^ and 4-2091^T^) exhibit distinct biosynthetic gene clusters (BGCs) responsible for secondary metabolite production. Strain A2^T^ contains two major BGCs: a T3PKS (Type III polyketide synthase) cluster and a terpene cluster, with the latter showing a 28% similarity to known carotenoid clusters. Strain A3E31^T^ also harbors a T3PKS cluster, alongside two other terpene clusters, one of which is closely related to carotenoids (28% similarity), while the other shows resemblance to flexirubin (41% similarity). On the other hand, strain PE311^T^ possesses three notable BGCs: a T3PKS cluster, an NRPS-like cluster combined with T1PKS (Type I polyketide synthase), and a terpene cluster that is similar to known carotenoid clusters (28% similarity). Strain 3-1016 ^T^ contains an NRPS cluster. Strain 4-2091^T^ also carries a T3PKS cluster with a similarity of 8% to NRP:B eta-lactam.(Table S7)

### Putative polysaccharide degradative capacity

The presence of degradative CAZymes and predicted PULs in each genome indicates the potential for polysaccharide degradation (Supplementary Table S8). Interestingly, the occurrence of PULs exhibited an exponential increase with the genome size, suggesting a correlation between genome size and polysaccharide-degrading capacity. Notably, marine isolates from algal species were annotated with a higher number of PULs, further underscoring their strong potential for polysaccharide degradation. These findings are consistent with previous studies that reported similar patterns of PUL distribution in marine organisms (50). However, as shown in Fig. 3, the distribution of sulfatases across the genomes did not exhibit a clear, consistent pattern, indicating that their presence may be more sporadic and potentially influenced by other factors. We observed a positive correlation between the numbers of CAZymes, sulfatases, and PULs and the different branches of the phylogenetic tree. Specifically, the branch positioned lower on the tree was annotated with a higher number of CAZymes, sulfatases, and PULs compared to the branches higher up the tree. However, exceptions to this trend are also evident, such as the case of A3E31^T^, which, despite being located on a more distantly related branch, is annotated with a relatively higher number of CAZymes, sulfatases, and PULs. This discrepancy may be linked to specific environmental factors or ecological conditions related to its habitat.

### Known and putative new laminarin PULs

Laminarins are β-1,3-linked glucans that are abundant as they act as storage compounds in brown algae and diatoms. Enzymes from the GH families 5, 8, 9, 16, 17, 30, 64, 81, and 157 are involved in laminarin degradation. The backbone is usually broken down by GH16_3 endo-glucanases (51). Just like the PUL structure (5) shown in the Fig.4, most strains can degrade laminarin and the PUL structure is relatively conserved. This reflects the fact that laminarin is one of the most abundant macromolecules in marine, as it acts as storage compound in brown macroalgae and diatoms. In the genus *Winogradskyella*, besides the common laminarin degradation pathway involving GH16, there is a distinct mode where GH65 family members exhibit endo-glucanase activity similar to that of GH16_3. This type of PUL is highly prevalent in *Winogradskyella*, accounting for 62 in all strains (e.g., 3-1016^T^). Therefore, future research may reveal the significant role of GH65-related enzymes in laminarin degradation, particularly in pathways that do not involve GH16, highlighting the potential of GH65 in glycoside hydrolysis.

**Fig. 4.:**
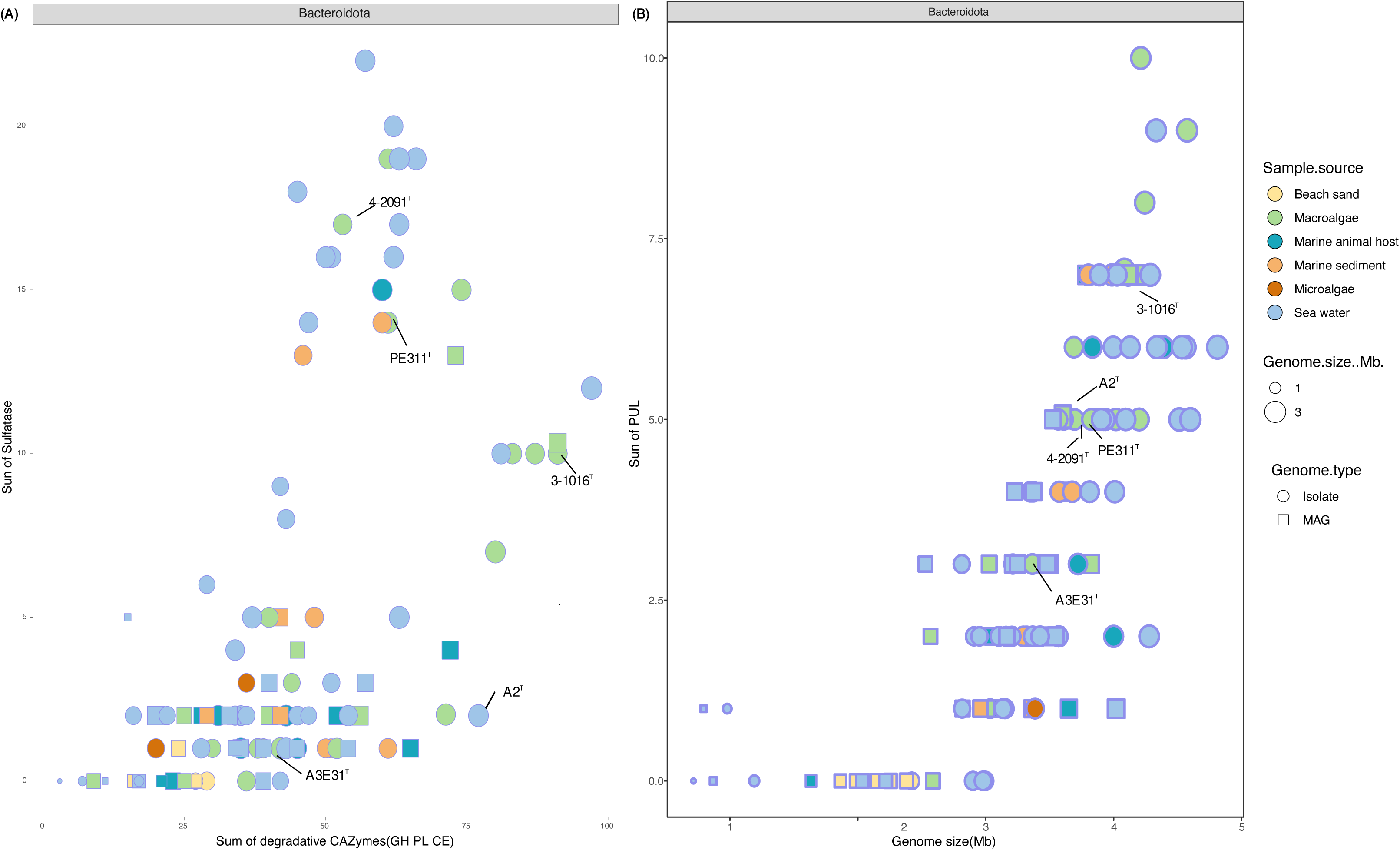
(A) Plot comparing the sum of degradative CAZymes (GH PL CE) (x-axis) and the sum of sulfatase enzymes (y-axis) for genomes and metagenome-assembled genomes (MAGs) within the *Winogradskyella*. (B) Plot comparing the sum of PULs (x-axis) and genome size (in Mb) (y-axis). Each point represents a different sample with circles for isolates and squares for MAGs. Colors represent different sample sources.

### Known and putative new Alginate

Alginates are linear co-polymers consisting of homopolymeric blocks of (1 → 4)-linked β-D-mannuronate and α-L-guluronate residues that are covalently linked in alternating sequences or blocks (52, 53). Alginates are anionic and bind sodium and calcium ions. Alginates were the second most frequently predicted PUL substrates in our dataset. They were predicted in 61 of the draft genomes, amounting to a total of 336 alginate-specific PULs. Alginate PULs encode PL6, 7, 14, 15, and 17 family alginate lyases.

Alginate, unlike laminarin, shows small variation across different strains in *Winogradskyella* genus. Sixty-four PULs (∼ 12%) in 48 genomes were predicted to be involved in alginate degradation, featuring three variants (Fig.5):

**Fig. 5.**
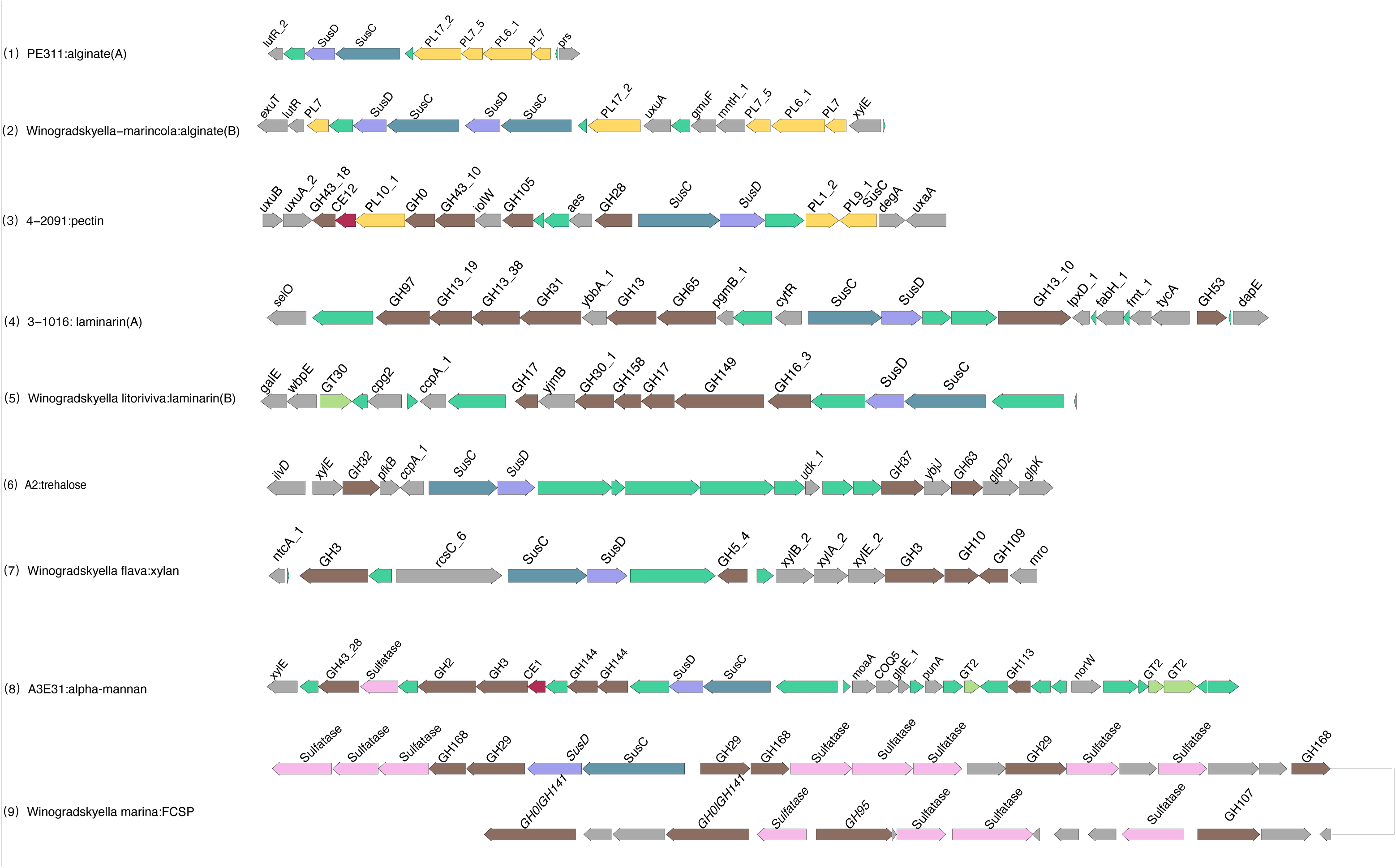
Selected PULs predicted to target substrates. Possible targets are (1) alginate A, (2) alginate B, (3) Pectin, (4) laminarin A, (5) laminarin B, (6) trehalose, (7) xylan and (8) alpha-mannan

Variant A is a highly conserved, corresponding PULs encode family PL6, 7, and 17 alginate lyases (Fig.6). This arrangement was first described in *Gramella forsetii* KT0803(53). For this species, it has been speculated that a cell surface-associated PL6 D-mannuronate-alginate lyase cleaves branched alginate polysaccharides into oligosaccharides (53, 54), which can be transported through the SusC-like TBDT into the periplasm. Here, the PL7 guluronate-specific alginate lyase may further cleave off glucose units (55), which are imported into the cytoplasm. But few type strains are of this PUL type. This may be due to the higher number of MAGs in the upper branch of the phylogenetic tree, where fewer PULs and CAZyme families are predicted compared to pure culture strains. Variants B PULs are likewise predicted to be capable of alginate degradation based on gene content but have not been described before. They feature an additional putative GH endo-b-1,2-glucosidase may be equivalent to the function of PL7. Variant C is the most widely distributed and most type species have this type of PUL. Interestingly, they possess two distinct SusC/SusD gene pairs. It suggests that they may possess a more diversified carbohydrate transport system. This increased diversity could enhance its ability to degrade a broader range of polysaccharides, may also enable them to process several substrates simultaneously, thereby improving its adaptability and survival in specific environments.

**Fig. 6.**
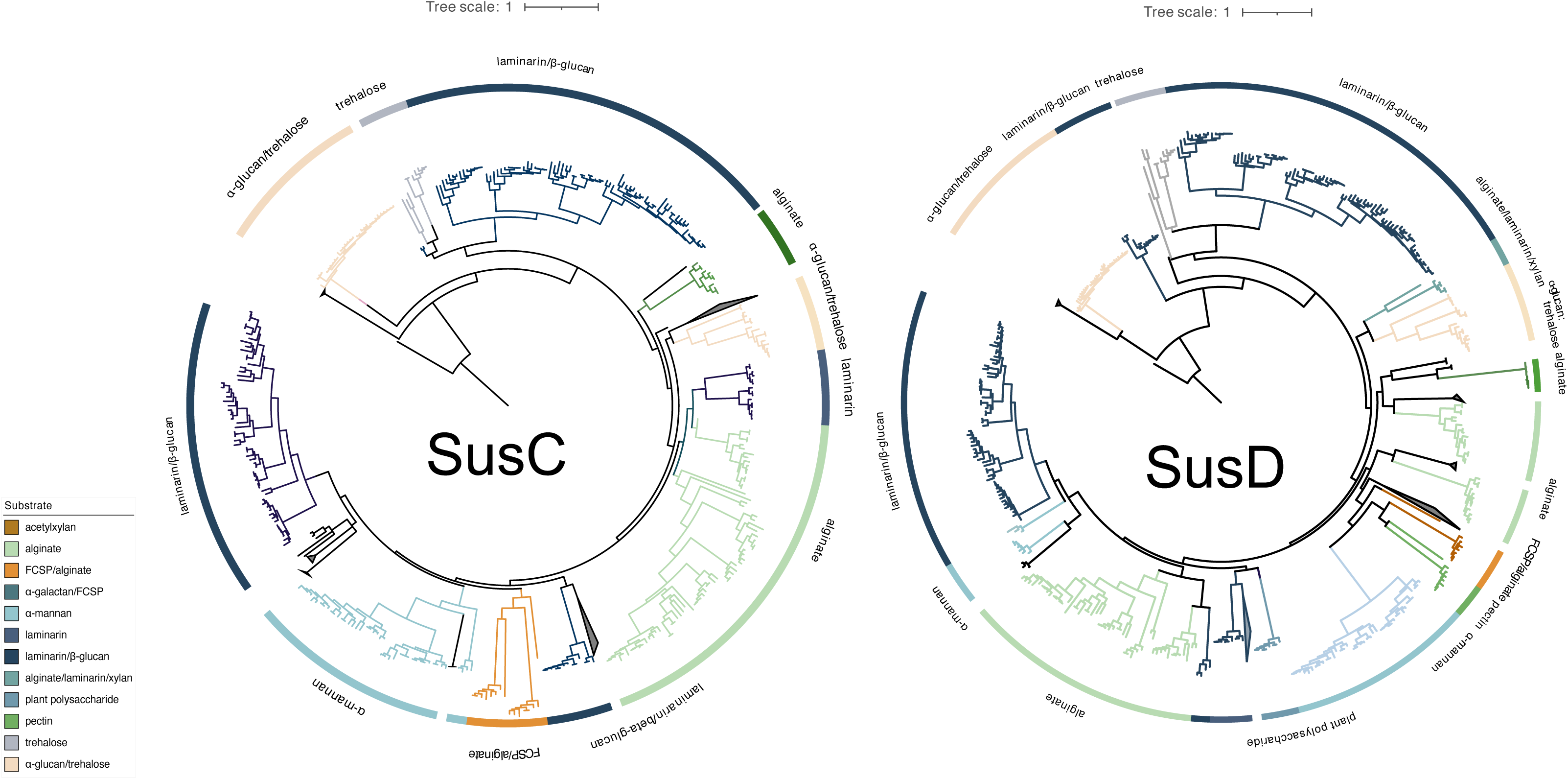
Trees of all PUL-associated SusC- (a) and SusD- (b) proteins of the *Winogradskyella* isolates showing functional, substrate-specific clustering. Protein sequences were aligned using the MAFFT G-INS-i algorithm and trees were calculated using FastTree 2.1.5 approximate-maximum likelihood. Substrate predictions are depicted in colors.

### Further substrates

Further possible substrates mainly include fucose-containing sulfated polysaccharides (FCSP), alpha-mannan, trehalose, and others (Fig.5). Fucoidans occur in brown macroalgae, such as *Fucus vesiculosus*, *Laminaria* spp. (kelp) and *Macrocystis* spp. (56). The main monomer is sulfated L-fucose, but the chemical composition of fucoidans is often complex and contains other monosaccharides (mannose, galactose, glucose, xylose, etc.), uronic acids, acetyl groups and even proteins (57). Known fucosidases are present in the GH29, 95, 107, 141 and 151 families, which is generally consistent with our findings. FCSPs are rich in sulfated esterases, thus requiring sulfatases, and the PULs we predicted are enriched with sulfatases. Predicted α-mannan-targeting PULs have been found to feature GH38, 76, 113, and 144 family CAZymes. They might be targeting α-glucomannans, such as glucuronomannan, a polysaccharide that has been reported for brown algae (58). Most α-mannan-targeting PULs code for GH113 family endo-mannosidases. In this study, 43 genomes featured a total of 488 PULs targeting mannose-rich substrates.

### Trees of SusC- and SusD-like proteins reveal substrate-specific clusters

We computed trees for all SusC- and SusD-like protein sequences of the 336 isolate PULs and obtained pronounced clusters for many of the predicted polysaccharide substrates (Fig. 7). For clarity, functionally heterogeneous or undefined clusters are depicted as gray triangles. Well-defined clusters in both trees included the structurally simple polysaccharides laminarin, α-1,4-glucans and alginate *et al*. This is visible in the trees by shorter and longer respective branch lengths. Some substrates formed multiple clusters, for example alginate and laminarin substrates. This might indicate either rather different alginate and laminarin or multiple ways of attack and uptake for a given class of alginate and laminarin. The topologies of the SusC- and SusD-like protein trees were notably congruent regarding branching patterns of the identified substrate-specific clusters. Alginate and laminarin cluster were located at a distinctly different position. SusC- and SusD-like proteins from the same PULs exhibited a strong tendency to occur in corresponding substrate-specific clusters in both trees. A considerable portion of the SusC and SusD sequences within the identified substrate-specific clusters showed similar patterns.

**Fig. 7.**
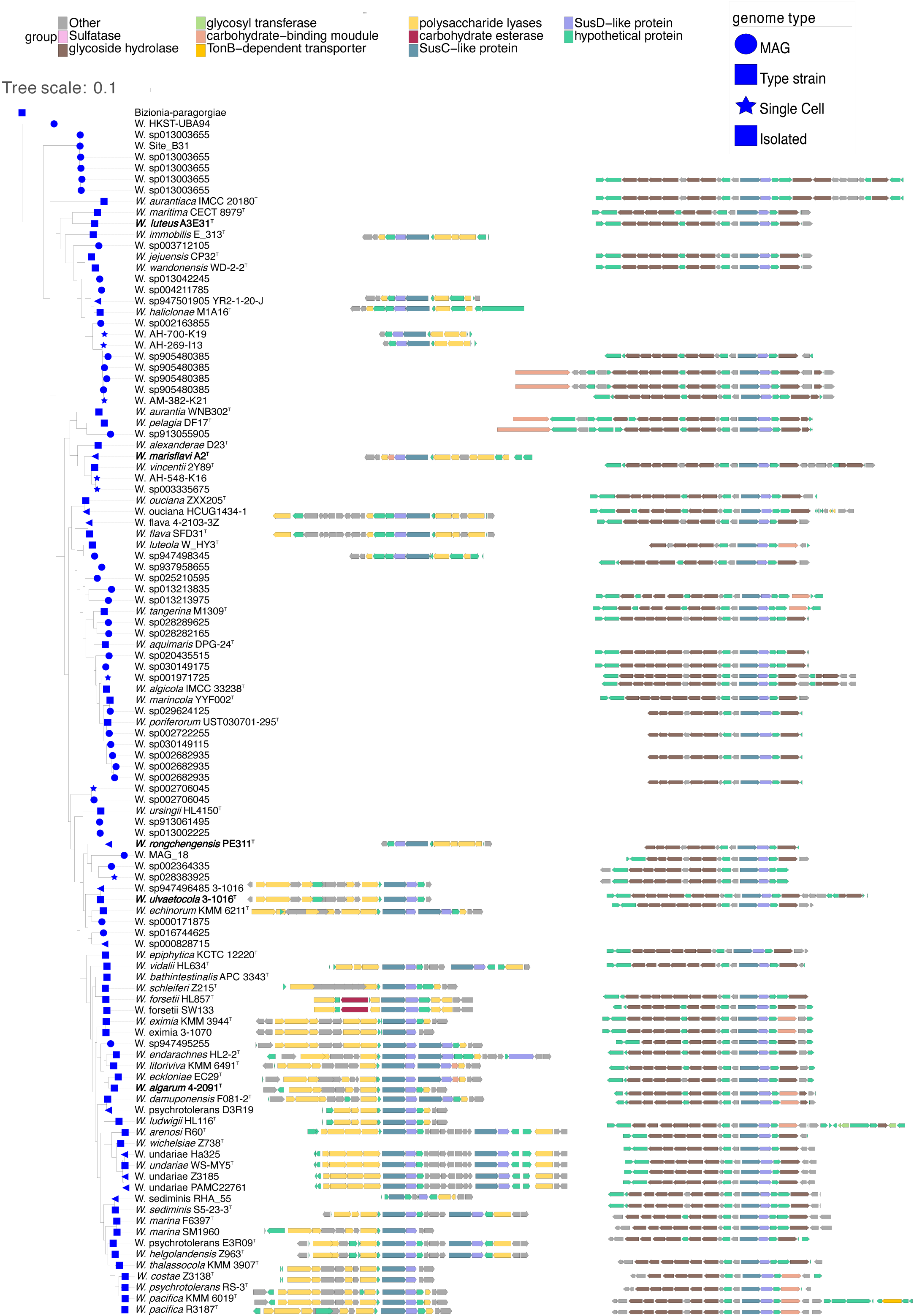
Distribution of PULs involved in the degradation of alginate and lamilarin across the phylogenetic tree. The left side represents alginate, and the right side shows lamilarin. Notably, lamilarin is more widely distributed within the *Winogradskyella* genus compared to alginate. Alginate degradation PULs can be further divided into two major branches, as illustrated in Fig. 5 PUL diagram (1) and (2).

### Conclusion and discission

Our research identified five novel *Winogradskyella* strains isolated from marine macroalgae and conducted a thorough phylogenetic, genomic and phenotypic analysis. Subsequently, by analyzing genome-wide, we conducted a basic comparative analysis of the genomic features, habitat, and metabolic potential of *Winogradskyella* strains. Furthermore, genome-based analysis revealed the production of secondary metabolites, including carotenoids and polyketides, by these *Winogradskyella* strains. These compounds are well known for their antimicrobial and anti-inflammatory properties, which suggest potential biotechnological applications in pharmaceuticals and environmental management. This discovery adds another layer to our understanding of the ecological roles of *Winogradskyella* and opens up possibilities for their use in industries focused on sustainable product development (59).

We then focused on analyzing the similarities and differences among *Winogradskyella* strains in terms of their polysaccharide degradation abilities for different polysaccharide substrates. We observed diverse polysaccharide degradation capacities among *Winogradskyella* strains, such as for laminarin and alginate. Through comparative analysis, we speculate the presence of new potential GH and PL families involved in the degradation of laminarin and alginate. Additionally, we observed a positive correlation between the genome size and the number of PULs, suggesting that strains with larger genomes may possess a stronger polysaccharide degradation capacity. This finding is consistent with the research by Jose et al. (2017), who noted a close relationship between genome size and ecological adaptation in marine *Flavobacteriaceae* strains. (60, 61)

We observed significant diversity in alginate degradation capabilities among the *Winogradskyella* genus, with no apparent direct correlation to their taxonomy. Even within the same genus, strains exhibited notably different PUL profiles and genome sizes. These results suggest that the composition of a species’ PUL repertoire is influenced more by its specific ecological niche than its phylogenetic lineage. This observation supports the hypothesis that PULs are subject to horizontal gene transfer within *Flavobacteriia*, facilitating adaptation to various ecological environments. (61, 62)

The PUL repertoires of the isolates reveal that common and structurally simple polysaccharides, such as laminarin, α-1,4-glucans, and alginate, are frequently targeted by conserved PULs. This suggests that maintaining the enzymatic machinery for degrading these substrates is advantageous for marine *Flavobacteriia*. The two alginate PUL variants we identified might reflect adaptation to different forms of alginate or serve as auxiliary modules that enhance the overall degradation process. Phylogenetic clustering of the SusC/D sequences of these variants further supports the notion that they interact with similar substrates.

A notable outcome of this study is the substrate-specific clustering of SusC- and SusD-like proteins. Our data indicate a strong tendency for these homologs to cluster in corresponding substrate-specific groups in phylogenetic trees, suggesting a coevolution of these proteins. This finding aligns with recent structural studies of *B. thetaiotaomicron* SusC/D complexes (63). We observed a more distinct clustering for simple, conserved polysaccharides, as opposed to heterogeneous substrates, which are cleaved at multiple sites to generate diverse oligosaccharides for uptake. This discrepancy reflects the complexity of substrate recognition in polysaccharide degradation, particularly for substrates that fall into broad categories like FCSPs or xylose-containing polysaccharides. These broad substrate classes may encompass chemically diverse compounds, and thus, further improvements in functional clustering are expected once more precise structural data on algal polysaccharides becomes available.

The PUL function predictions in this study rely heavily on bioinformatic analyses based on sequence similarities, which, while useful, cannot achieve the precision of more laboratory techniques. Given the limited knowledge on polysaccharides from marine macroalgae, some predictions may be inaccurate. However, the holistic approach used to analyze PUL profiles across a wide array of isolates from a single habitat allows for the identification of common PULs that are likely to be biologically significant. These recurring PULs represent crucial targets for experimental studies and provide a solid foundation for developing hypotheses about potential polysaccharide substrates. Despite its limitations, this approach offers a novel framework for identifying environmentally relevant polysaccharide substrates that remain challenging to characterize through direct chemical analysis.

Future studies should combine genomic analyses with metabolic pathway investigations to enhance our understanding of the ecological roles of these strains and explore their potential applications in marine ecosystems.

## Abbreviations

KCTC: Korea Collection for Type Culture
MCCC: Marine Culture Collection of China
MA: marine agar
NCBI: National Centre for Biotechnology Information
ANI: average nucleotide identity
GGDC: Genome-to-Genome Distance Calculator
NJ: Neighbor-Joining
ML: Maximum-Likelihood
ME: Minimum-Evolution
PE: phosphatidylethanolamine
L: unidentified lipids
AL: unidentified aminolipid
PG: phosphatidylglycerol
MK-6: Menaquinone 6
GH: glycoside hydrolases
GT: glycosyl transferases
CE: carbohydrate esterases
CBM: carbohydrate-binding modules
AA: auxiliary activities
PL: polysaccharide lyases
CAZyme: carbohydrate-active enzyme
PUL: polysaccharide utilization loci
CGC: CAZyme-rich gene clusters
BGC: biosynthetic gene clusters
NRPS: non-ribosomal peptide synthetases.

## Data availability

The GenBank accession number for the 16S rRNA gene sequence of strain A3E31^T^, PE311^T^, A2^T^, 4-1091^T^ and 3-1016^T^ is PQ357589, PQ357496, PQ421146, PQ609710 and PV241781 respectively. The draft genome of strain A3E31^T^, PE311^T^, A2^T^, 4-2091^T^ and 3-1016^T^ had been deposited in GenBank under the accession number JBHNWH000000000, JBHZTH000000000, JBKQXM000000000, JBJKOD000000000 and CANMCZ000000000 respectively. BioSample: SAMEA112156868, SAMN40544020; BioProject: PRJEB57783. Sequences are available from the European Nucleotide Archive under accessions PRJEB50838 (metagenomes and MAGs), and PRJEB57783 (genomes of cultured bacteria).

## Acknowledgements

This work was supported by the Science Foundation for Youths of Shandong Province (ZR2023QC197), the Science and Technology Fundamental Resources Investigation Program (Grant number 2022FY101100) and the National Natural Science Foundation of China (92351301). The scanning electron microscopy was supported by the Physical-Chemical Materials Analytical and Testing Center of Shandong University at Weihai.

## Declarations

Conflicts of interest Authors declare that there was no conflict of interest. This article does not contain any studies with animals performed by any of the authors.

## Ethics approval

Not applicable.

## Consent to participate

Informed consent was obtained from all individual participants included in the study.

## Consent for publication

All authors have agreed to it being published.

